# Building Functional Connectivity Neuromarkers of Behavioral Self-Regulation in Children with and without Autism Spectrum Disorder

**DOI:** 10.1101/635292

**Authors:** Christiane S. Rohr, Shanty Kamal, Signe Bray

**Affiliations:** Child and Adolescent Imaging Research Program, The University of Calgary, Calgary, Alberta, Canada; Alberta Children’s Hospital Research Institute, The University of Calgary, Calgary, Alberta, Canada; Mathison Centre for Mental Health, The University of Calgary, Calgary, Alberta, Canada; Hotchkiss Brain Institute, The University of Calgary, Calgary, Alberta, Canada; Department of Radiology, Cumming School of Medicine, The University of Calgary, Calgary, Alberta, Canada; Department of Paediatrics, Cumming School of Medicine, The University of Calgary, Calgary, Alberta, Canada

**Keywords:** shifting, cognitive flexibility, inhibition, connectome predictive modelling, fingerprinting

## Abstract

Children with Autism Spectrum Disorder (ASD) are known to struggle with behavioral self-regulation, which associates with greater daily-life challenges and an increased risk for psychiatric comorbidities. Despite these negative outcomes, little is known about the neural expression of behavioral regulation in children with and without ASD. Here, we examined whole-brain linear associations between brain functional correlations (FC) and behavioral regulation through connectome predictive modelling (CPM), a data-driven protocol for developing predictive models of brain–behavior relationships from data, assessing ‘neuromarkers’ using cross-validation. Using data from two sites of the ABIDE II dataset comprising 276 children with and without ASD (8-13 years), we identified functional brain networks whose FC predicted individual differences in two, of three, behavioral regulation subdomains. These distributed network models predicted novel individuals’ inhibition and shifting from resting-state FC data both in a leave-one-out, as well as split halves, cross-validation. We observed commonalities and differences in the functional networks associating with these subdomains, with inhibition relying on more posterior networks, shifting relying on more anterior networks, and both involving regions of the DMN. Our findings present a substantial addition to our knowledge on the neural expressions of inhibition and shifting across the spectrum of children with and without ASD, demonstrating the utility of this trans-diagnostic modelling approach. Given the numerous cognitive and behavioral issues that can be quantified dimensionally in neurodevelopmental disorders, further refinement of whole-brain neuromarker techniques may thus pave a way for functional neuroimaging to meaningfully contribute to individualized medicine.

## 1 INTRODUCTION

Children with Autism Spectrum Disorder (ASD) are known to struggle with behavioral self-regulation and exhibit problems with executive functions more generally [1–3]. Executive function difficulties are theoretically and experimentally linked to the diagnostic symptoms of children with ASD, such as: difficulty with normed social interactions and communication; circumscribed interests and repetitive behaviors; and both internal (anxiety, depression) and external (aggression, hyperactivity) emotional struggles [3–13]. Yet, little is known about the brain correlates of behavioral regulation in ASD.

Individuals with ASD have trouble in multiple aspects of behavioral regulation subdomains, including inhibitory control processes, cognitive flexibility, and emotional control processes [11, 14–16]. Inhibitory control is the ability to suppress interfering distractions and prepotent motor responses [11, 14, 17]. Studies indicate that both reduced cognitive and motor inhibition are present in ASD throughout the lifespan [2, 11, 18, 19]. Another integral facet of behavioral regulation is cognitive flexibility, which is typically measured using set-shifting or the readiness with which one can switch from one task or mindset to another [14, 20, 21]. Reduced cognitive flexibility is readily observed in the rigidity of ASD behaviors [22, 23]. Finally, emotional control, also termed ‘cognitive control of emotion’ or emotion regulation, is the processes by which we influence which emotions we have, when we have them, and how we experience and express them [24, 25], is also known to be hampered in ASD [3, 26, 27].

The brain networks implicated in these three subdomains of behavioral regulation are heavily intertwined. Common pathways - largely identified through task-based fMRI studies - most prominently revolve around areas in the prefrontal cortex (PFC), such as the ventrolateral PFC (vlPFC), the dorsolateral PFC (dlPFC), ventromedial and dorsomedial PFC (vmPFC/dmPFC), as well as the anterior cingulate cortex (ACC). For example, data suggests that the vlPFC supports reflexive reorienting, motor inhibition, and action updating [28], the selection of the most efficacious response set when confronted with a task requiring various possible responses [29, 30], and deliberately increasing or decreasing negative affect [31]. Enhanced dlPFC activity and also enhanced activity in medial prefrontal structures is often observed, in conflict paradigms, which requires inhibitory control of prepotent but incorrect responses and set-shifting to reframe the problem [32]; such paradigms include emotional conflict paradigms where the preprotent responses that need to be controlled are emotional [33–35]. In addition, areas in the parietal and temporal lobes, such as the inferior parietal lobule (IPL), the superior temporal gyrus (STS) and the temporal pole, as well as limbic structures, most notably the insula and amygdala, are known to play important roles in cognitive flexibility [36] and emotion control [33, 37–39].

Despite a dominant focus on prefrontal brain regions in the behavioral regulation literature, distributed regions and their interactions have also been implicated. For example, changes in inter-regional synchrony, i.e. functional connectivity or correlation (FC), during emotional control processes have been observed between IPL and vmPFC, and STS and dmPFC [33], as well as between parietal structures and the amygdala (Ferri). In addition, meta-analytic evidence [36] lends further support to the involvement of a distributed network including parietal and temporal areas and the importance of (FC). Importantly, FC can be measured both during behavioral regulation tasks, as in the literature noted above, as well as at rest, where it is thought to reflect intrinsic functional brain organization that is reflective of an individual’s attributes, including cognitive and behavioral traits [38, 40, 41]. In ASD, distributed network patterns have been associated with social symptoms [42], but less is known about distributed FC in relation to behavioral regulation.

It should be noted that while the neural substrates of executive functioning and behavioral regulation have been extensively studied in neurotypical adults and adolescents/young adults with affective or developmental disorders, less is known about the neural expression of behavioral regulation in children and children with ASD [8, 43]. The neural signatures of ASD have been elusive due to heterogeneity in the condition, so taking a dimensional approach to look at specific features may be more promising [42, 44, 45]. In particular, whole-brain FC signatures that define aspects of behavioral regulation in ASD have been elusive, despite the widespread repercussions of suboptimal behavioural regulation throughout development and into adulthood for children with ASD, and the enormous potential of whole-brain FC profiles as ‘neuromarkers’ for diagnosis and individually tailored treatment.

Here we use connectome predictive modelling (CPM), a data-driven protocol for developing predictive models of brain–behavior relationships from FC data using cross-validation [46], to examine whole-brain linear associations of behavioral regulation and their utility as ‘neuromarkers’. We hypothesize (a) that FC models of behavioural regulation can be built across an aggregate sample of children with and without ASD, (b) that frontoparietal, limbic and default mode networks underlie behavioral regulation, and (c) that neuromarkers built in a subset of the sample can then be used to predict behavioral regulation scores in another, unseen subset of children.

## 2 METHODS

### 2.1 Participants

For this study we used two datasets from the Autism Brain Imaging Data Exchange II (ABIDE-II) database [47]. These sites were chosen because they included Behavior Rating Inventory of Executive Function (BRIEF [16]) behavioural regulation scores and resting state fMRI data from children aged 8-13 years with and without ASD. Specifically, we analyzed the datasets collected at Georgetown University (GU) and the Kennedy Krieger Institute (KK), which are publicly available under fcon_1000.projects.nitrc.org/indi/abide/abide_II.html. At both sites, children were introduced to an MRI simulator first and given the opportunity to familiarize themselves with the experience of undergoing an MRI scan with their eyes open. Potential participants were excluded if they had a history of neurological or psychiatric disorders (the latter only in the typically developing (TD) group), contraindications for MRI, or if they had other medical problems that prevented participation. Nine participants did not have behavioral data on file and were excluded from the analysis. Participants’ data were further evaluated for outliers in behavioral measures, high inconsistency scores on the BRIEF or excessive motion on the fMRI scans. Specifically, for the behavioral measures, outliers were defined as > 3 SD from the mean. One participant was excluded due to this criterion in the subdomain of emotion control and shifting, and 6 in the subdomain of inhibition. An additional participant was excluded due to this criterion in analyses that involved the social responsiveness scale (SRS). Five participants were excluded because of inconsistency scores >7. Twenty-six participants had excessive head motion on their fMRI scan (>4 mm maximum absolute displacement). The final samples and participant characteristics are given in Table 1.

**Table 1.** Participant characteristics in the GU site, the KK site and the combined sample. Means and standard deviations (in brackets) are given for the total samples comprised of both TD and ASD participants, which were used to build the models, as well as for TD and ASD participants separately. Motion (mm) refers to the absolute maximum displacement at any timepoint in the resting-state fMRI scan prior to motion mitigation and denoising procedures. n=number of participants; m=male; f=female; L=left-handed; A=ambidextrous; R=right-handed; IQ=Intelligence Quotient; FIQ=full scale IQ; VIQ=verbal IQ; PIQ=performance IQ; SRS=Social Responsiveness Scale (total score). SRS, inhibition, shifting and emotion control are given as T scores. * denotes deviating numbers in the Inhibition models, for which an additional six outliers (>3 SD in score; all ASD participants) were removed. ** SRS scores were not available for some participants in the KK site (140 out of 145 TD and 47 out of 49 ASD); 1 participant with ASD from the GU site was removed as an outlier in SRS scores.

|  | GU Site |  |  | KK Site |  |  | Combined Sample |  |  |
| --- | --- | --- | --- | --- | --- | --- | --- | --- | --- |
|  | Total | TD | ASD | Total | TD | ASD | Total | TD | ASD |
| n | 82 | 46 | 36 | 194 | 145 | 49 | 276 | 191 | 85 |
| Age | 10.8 (1.6) | 10.5 (1.7) | 11.2 (1.4) | 10.3 (1.3) | 10.3 (1.2) | 10.3 (1.5) | 10.5 (1.4) | 10.4 (1.3) | 10.7 (1.5) |
| Sex (m/f) | 54/28 | 23/23 | 31/5 | 128/66 | 93/52 | 35/14 | 182/94 | 116/75 | 66/19 |
| Handedness (L/A/R) | 9/0/73 | 3/0/43 | 6/0/30 | 12/16/166 | 10/10/125 | 2/6/41 | 21/16/239 | 12/10/168 | 8/6/71 |
| Motion | 1.1 (0.7) | 1.1 (0.7) | 1 (0.7) | 1.4 (0.9) | 1.2 (0.8) | 1.8 (0.9) | 1.3 (0.9) | 1.2 (0.8) | 1.5 (0.9) |
| FIQ | 120.2 (14.2) | 121.7 (14.2) | 118.3 (14.3) | 111.9 (12.7) | 114.6 (10.4) | 103.7 (15.5) | 114.3 (13.7) | 116.3 (11.8) | 109.8 (16.6) |
| VIQ | 121.8 (14.9) | 121.8 (15.5) | 121.7 (14.4) | 116 (13.7) | 118.2 (11.8) | 109.4 (16.9) | 117.7 (14.3) | 119.1 (12.8) | 114.6 (16.8) |
| PIQ | 115.9 (13.7) | 116.9 (13.4) | 114.1 (14.4) | 109.4 (12.8) | 110.8 (12.1) | 105.4 (14.3) | 111.1 (13.4) | 112.2 (12.6) | 108.4 (14.8) |
| SRS** | 56.3 (17.2) | 43.9 (6.4) | 72.6 (12.5) | 51.5 (16.1) | 43.2 (5.3) | 76.4 (10.7) | 53 (16.5) | 43.4 (5.6) | 74.8 (11.5) |
| Inhibition* | 50.1 (9.8) | 44.9 (6.3) | 56.8 (9.5) | 47.9 (10) | 43.8 (6.1) | 61.7 (8.4) | 48.6 (10) | 44.1 (6.1) | 59.4 (9.2) |
| Shifting | 53.8 (14.6) | 44.6 (6.8) | 65.5 (13.5) | 49.7 (13.9) | 43.4 (7.1) | 68.5 (12) | 50.9 (14.2) | 43.7 (7) | 67.2 (12.7) |
| Emotion Control | 51.5 (11.7) | 45.2 (7.7) | 59.6 (11.1) | 47.5 (11) | 43.5 (7.1) | 59.4 (11.8) | 48.7 (11.3) | 43.9 (7.3) | 59.5 (11.5) |

### 2.2 Cognitive and Behavioral Assessment

The cognitive and behavioral data used for analysis included IQ, handedness, ASD symptoms and behavioral regulation scores. IQ was measured using the Wechsler Intelligence Scale for Children – Fourth or Fifth Edition (WISC IV [48]; WISC-V [49]) across the KK site, while in the GU site it was assessed either using the WISC-IV or the Wechsler Abbreviated Scale of Intelligence (WASI [50]). Handedness was determined using the Edinburgh Handedness Inventory [51] in the KK site; in the GU site handedness was obtained by self- and parent-report. As an estimate of ASD symptoms, social responsiveness was measured using the Social Responsiveness Scale (SRS), Edition 1, version 1 [52] in the GU site and either in Edition 1, version 1 or Edition 2, version 1 [53] in the KK site. Behavioral regulation in the datasets was assessed with the BRIEF [16], a parent assessment of executive function behaviors for children aged 5-18 years. BRIEF subscales provide measures of three domains of behavioral regulation, which are labelled “inhibit”, “shift”, and “emotional control”. The “inhibit” subscale assesses the ability to resist impulses and to stop one’s own behavior” (sample item: “acts wilder or sillier than others in groups (birthday parties, recess)”). The “shift” subscale assesses the ability to move freely from one situation, activity, or problem to another; to tolerate change, and to switch or alternate attention (sample item: “resists or has trouble accepting a different way to solve a problem with schoolwork, friends, chores, etc.”). Finally, the “emotional control” subscale assesses the ability to regulate emotional responses appropriately (sample item: “overreacts to small problems”). T-scores for both the SRS and the BRIEF were used for analysis.

### 2.3 Analysis of Cognitive and Behavioral Measures

To assess differences in characteristics (demographics, cognitive abilities and outcomes of interest) between children with and without ASD, as well as between groups within and across the two sites, t-tests were computed. Further, to assess the relationship between behavioral regulation scores, ASD symptom scores and IQ, Pearson correlations were computed. One-way ANOVAs were used to assess potential differences in behavioral regulation scores, ASD symptom scores and IQ in relation to handedness. Behavioral analyses were carried out using SPSS 22 (Chicago, IL).

### 2.4 MRI Data Acquisition Parameters

Data were acquired on a 3T Siemens Magenotom TrioTim at the GU site, and on a 3T Philips Achieva at the KK site. Children were instructed to keep their eyes open at both sites. Functional images were acquired using a gradient-echo EPI sequence in 43 axial slices (154 volumes, TR=2000 ms, TE=30 ms, FA=90, matrix size 64×64, voxel size 3×3×3 mm^3^; duration: 5.14 minutes) at the GU site, and in 47 axial slices (128 volumes, TR=2500 ms, TE=30 ms, FA=75, matrix size 96×96, voxel size 2.67×2.67×3 mm^3^; duration: 5.3 minutes) at the KK site. Anatomical scans were acquired using a T1-weighted MPRAGE sequence (GU: TR=2530 ms, TE=3.5 ms, FA=7, voxel size 1×1×1 mm^3^; KK: TR=8.2 ms, TE=3.7 ms, FA=8, voxel size 1×1×1 mm^3^).

### 2.5 MRI Data Preprocessing

Data preprocessing was done using functions from FSL [54] and AFNI [55]; the specific functions are denoted in brackets. Anatomical data was deobliqued (3drefit), oriented into FSL space (RPI) (3dresample) and skull-stripped (3dSkullStrip and 3dcalc). Functional data was also first deobliqued (3drefit) and oriented into FSL space (RPI) (3dresample). The pipeline further consisted of motion correction (MCFLIRT), skull-stripping (3dAutomask and 3dcalc), spatial smoothing (6 mm Gaussian kernel full-width at half-maximum) (fslmaths), grand-mean scaling (fslmaths), registration to the participants anatomical (FLIRT), and normalization to the McConnell Brain Imaging Center NIHPD asymmetrical (natural) pediatric template optimized for ages 7.5 to 13.5 years [56] (FLIRT), followed by normalization to 2×2×2 mm MNI152 standard space (FLIRT).

### 2.6 Head-motion and Physiological Confound Mitigation Procedure

We used a four-step process to address motion and physiological confounds in the data. First, we used motion estimates derived from the preprocessing in order to exclude participants with excessive head motion; scans were excluded if they exhibited > 4 mm maximum absolute displacement. Second, on the participants who were retained for analysis, we used AROMA, an ICA-based cleaning method [57], which has recently been shown to be most effective in mitigating the impact of head motion, and allows for the retention of the remaining ‘true’ neural signal within an affected volume [58]. AROMA is an automated procedure that uses a small but robust set of theoretically motivated temporal and spatial features (timeseries and power spectrum) to distinguish between ‘real’ neural signals and motion artifacts. We chose a conservative threshold (‘aggressive’) in order to decrease the chance of false positives. In other words, more components are removed as this threshold is more conservative about what is retained. Noise components identified by AROMA were removed from the data. Third, images were de-noised by regressing out the six motion parameters, as well as signal from white matter, cerebral spinal fluid and the global signal, as well as their first-order derivatives [59]. While there is currently no gold standard [60] regarding the removal of the global signal, we chose to remove it based on recent evidence that it relates strongly to respiratory and other motion-induced signals, which persist through common denoising approaches including ICA and models that attempt to approximate respiratory variance [61]. Motion (defined as each participants’s absolute maximum displacement) was substantially reduced following this procedure (before: 1.28mm ± 0.85mm; after: 0.05mm ± 0.07mm). As a final step, as described in more detail below, head motion was incorporated into models by removing connections that remained significantly (p<0.05) associated with z-scored motion before cleaning in a Pearson’s correlation [62].

### 2.7 Connectome-Predictive Modelling

To elucidate how behavioral regulation skills are reflected in children’s whole-brain FC profiles (or ‘connectomes’), and how they vary across children with and without ASD, we utilized a protocol termed Connectome Predictive Modelling (CPM). CPM is an algorithm for building predictive models based on participants’ FC matrices, and for testing these models using cross-validation of novel data. Scripts are written in MATLAB and are freely available at www.nitrc.org/projects/bioimagesuite. The CPM protocol is described in detail in [46] and has previously been applied to pediatric data sets including data from the ABIDE sample [42, 62–64]. We followed the CPM protocol [46], as well as recent recommendations for predictive modelling [65], in calculating each participant’s FC profile, building models of behavioral regulation, and in running the following analyses: (1) a leave-one-out cross-validation to evaluate the potential of models to predict an unseen participant’s score, where N-1 participants are used to build the predictive model and the model is subsequently tested on the left-out participant; (2) a split-halves prediction where all available data was randomly split and models built in the first half were used to predict individuals in the second half and vice versa; (3) a site-to-site prediction where models built in the GU site were used to predict individuals in KK site and vice versa. We describe how we calculated FC matrices, built the models and ran these analyses in the following sections.

#### 2.7.1 Calculation of FC profiles

A functionally defined atlas, consisting of 268 cortical and subcortical regions-of-interest (‘nodes’) that cover the whole brain [66], was used to define ROIs. For each child, we extracted the timecourse of each node by taking the mean across voxels and a 268×268 connectivity matrix was calculated between timecourses of node pairs using Pearson’s correlation followed by Fisher’s Z transformation. Thus, each connection (or ‘edge’) in the matrix represents the strength of FC between two nodes, and the matrix as a whole represents a child’s FC profile or functional connectome.

#### 2.7.2 Building FC Models of Behavioral Regulation

Models were built relating z-scored behavioral regulation subscales (emotional control, shift, inhibit) to FC matrices across participants with and without ASD from both sites. Prior to modeling, effects of motion and site were eliminated from participants’ FC profiles by masking out connections that were significantly (p<0.05) associated with motion in a Pearson correlation or different between sites in a t-test. 6915 out of a possible 35778 (=268×267, adjusted for the diagonal, divided by 2, because matrices are symmetric) nodes were eliminated at this step due to motion, leaving 28863 valid nodes in the matrix; accounting for site brought this number down to 23100. For model building, each edge in the matrix was correlated with the behavioral regulation measures (again in a Pearson’s correlation), and only significantly correlated edges (p<0.01) were selected and retained. These selected edges were first separated into positively and negatively associated edges based on the direction of the correlation, as they may be interpreted differently in terms of their functional roles, and then summed for each participant, yielding a single summary FC value per participant for each of the positive and negative edge models. In other words, each participant FC had two summary FC values, one for a network that positively associated with behavioral regulation and one for a network that negatively associated with behavioral regulation. Finally, a predictive model was built that fits a linear regression between each participants’ summary FC value and the behavioral regulation variable of interest [46].

#### 2.7.3 Cross-Validation: Leave-One-Out Prediction

To evaluate the potential of models to predict an unseen participant’s score, one participant was removed from the dataset and the remaining participants (N-1) are used to build the predictive model. The left-out-participant’s score was predicted based on the N-1 sample’s fit of the linear regression model, and this step is repeated in an iterative manner with a different participant left out in each iteration. Spearman’s r_s_ was used to evaluate model performance i.e. comparing actual to predicted scores because it is less sensitive to the effect of outliers than Pearson’s r and because CPM predictions are best considered relative rather than absolute [62, 63]. Only models that showed a significant correlation at p<0.05 between observed and predicted scores at this step were subjected to follow-up testing.

#### 2.7.4. Evaluation of the Predictive Model

The predictive potential is assessed by comparison of the predicted and observed scores in the full model, and statistical significance is assessed using permutation testing (5000 iterations). Permutation (i.e., randomization) testing was used to assess significance because the assumption underlying the standard r-to-p conversion that was employed in the leave-one-out cross-validation (see above) is violated: folds are not independent and thus the number of DOF is over-estimated [62, 63, 65]. To perform permutation testing, we randomly shuffled participants’ behavioral scores 5000 times and ran these shuffled values through our prediction pipeline to generate null distributions. P-values associated with each model were based on the corresponding null distribution with the formula p = (1 + the number of permutation r_s_ values greater than or equal to the observed r_s_ value)/5001 [62]. In other words, the p-value of the permutation test is the proportion of sampled permutations that are greater than the true prediction correlation [46].

#### 2.7.5 Cross-Validation: Split Halves Prediction

As a further test of the models, data from both sites were randomly split while retaining the same number of TD and ASD participants and the same number of participants from each site. Models were built in the first half and used to predict individuals in the second half and vice versa. Participant characteristics for the two split halves are given in Table 2.

**Table 2.** Participant characteristics in the two split halves samples. Means and standard deviations (in brackets) are given for the total samples comprised of both TD and ASD participants, which were used to build the models, as well as for TD and ASD participants separately. Motion (mm) refers to the absolute maximum displacement at any timepoint in the resting-state fMRI scan prior to motion mitigation and denoising procedures. n=number of participants; m=male; f=female; L=left-handed; A=ambidextrous; R=right-handed; IQ=Intelligence Quotient; FIQ=full scale IQ; VIQ=verbal IQ; PIQ=performance IQ; SRS=Social Responsiveness Scale. SRS, inhibition, shifting and emotion control are given as T scores. *denotes deviating numbers in the Inhibition models, for which an additional six outliers (>3 SD in score; all ASD participants) were removed. **SRS scores were not available for seven participants in Split Half 1; 1 participant with ASD was removed as an outlier from Split Half 1.

|  | Split Half 1 |  |  | Split Half 2 |  |  |
| --- | --- | --- | --- | --- | --- | --- |
|  | Total | TD | ASD | Total | TD | ASD |
| n | 138 (135*) | 95 | 42 (39*) | 138 (135*) | 96 | 49 (43*) |
| Age | 10.6 (1.4) | 10.5 (1.4) | 10.6 (1.5) | 10.3 (1.3) | 10.2 (1.2) | 10.3 (1.5) |
| Sex (m/f) | 84/54 | 53/42 | 35/7 | 98/40 | 63/33 | 35/14 |
| Handedness (L/A/R) | 8/7/123 | 5/5/85 | 3/2/38 | 13/9/116 | 8/5/83 | 5/4/33 |
| Motion | 1.2 (0.8) | 1.1 (0.8) | 1.6 (0.9) | 1.3 (0.9) | 1.2 (0.8) | 1.8 (0.9) |
| FIQ | 113.6 (12.6) | 114.7 (11.3) | 108.7 (18.1) | 115.1 (14.7) | 117.8 (12.1) | 103.7 (15.5) |
| VIQ | 117.6 (14.2) | 117.8 (12.9) | 112.1 (16.7) | 117.8 (14.4) | 120.3 (12.5) | 109.4 (16.9) |
| PIQ | 110.4 (12.4) | 111 (12.3) | 107.7 (16.9) | 111.9 (14.3) | 113.5 (12.9) | 105.4 (14.3) |
| SRS** | 52.9 (17.6) | 42.7 (5.8) | 76.5 (12.1) | 53 (15.4) | 44.1 (5.4) | 73.1 (10.9) |
| Inhibition* | 48.3 (9.9) | 43.5 (5.4) | 59.2 (9.7) | 48.9 (10.1) | 44.7 (6.8) | 61.7 (8.4) |
| Shifting | 50.4 (14.3) | 43.1 (6.9) | 67.8 (12.4) | 51.5 (14.1) | 44.3 (7.1) | 68.5 (12) |
| Emotion Control | 48.6 (11.4) | 43.1 (6.1) | 58.3 (11.99) | 48.9 (11.4) | 44.7 (8.3) | 59.4 (11.8) |

#### 2.7.6 Cross-Validation: From Site 1 to Site 2 Prediction

To evaluate the potential of models built in one site to predict an unseen participant’s score from the other site, models were built in the GU site and used to predict individuals in the KK site and vice versa.

#### 2.7.7 Model Specificity to Behavioral Regulation

To evaluate whether our models capture behavioral regulation dimensionally or are driven by the categorical difference in scores due to ASD diagnoses, we examined the relationship between predicted and observed scores for TD children and children with ASD separately. We further examined (a) whether our models could be driven by motion or IQ, (b) how they relate to the core symptoms of ASD, and (c) whether models built for one domain of behavioral regulation were specific to that domain by computing cross-correlations between (c1) predicted scores and SRS scores, and (c2) predicted and observed scores of different behavioral regulation domains.

## 3 RESULTS

### 3.1 Sample Characteristics

Characteristics for all samples (GU site, KK site, combined sites; split half 1, split half 2) are given in Tables 1 and 2. Scores for all three subscales of behavioral regulation were significantly higher in children with ASD than for TD children in all analyzed samples, reflecting greater challenges with inhibition, shifting, and emotional control (Table 3). We further observed significantly higher scores on social responsiveness, reflective of greater ASD symptoms, and several significant differences within and across some of the samples in age, IQ, sex and head motion (Table 2). As expected, the three subscales of behavioral regulation exhibited correlations with each other as well as to social responsiveness and, to a lesser degree, IQ and head motion (Table 4). There were no significant differences in handedness between TD children and children with ASD in any of the samples.

**Table 3.** Significance of group differences between TD children and children with ASD. Scores for all three subscales of behavioral regulation were significantly higher in children ASD than for TD children in all analyzed samples, reflecting more issues with inhibition, shifting, and emotional control. We observed that in some, but not all samples, ASD children moved more and had lower IQs. In addition, akin to the ratio in the general population, ASD children tended to be predominantly male. Results are given as p-values of independent sample t-tests, which are adjusted in cases of unequal variance as assessed through Levene’s test. Motion (in mm) refers to the absolute maximum displacement at any timepoint in the resting-state fMRI scan before and after motion mitigation and denoising procedures. SRS=Social Responsiveness Scale (total score); IQ=Intelligence Quotient; FIQ=full scale IQ; VIQ=verbal IQ; PIQ=performance IQ.

| ASD > TD | GU Site | KK Site | Combined Sample | Split Half 1 | Split Half 2 |
| --- | --- | --- | --- | --- | --- |
| Age | 0.045 | - | - | - | - |
| Males | < 0.0001 | - | 0.004 | - | 0.022 |
| Motion (before) | - | - | 0.007 | - | 0.022 |
| Motion (after) | < 0.0001 | - | - | - | - |
| SRS | < 0.0001 | < 0.0001 | < 0.0001 | < 0.0001 | < 0.0001 |
| Inhibition | < 0.0001 | < 0.0001 | < 0.0001 | < 0.0001 | < 0.0001 |
| Shifting | < 0.0001 | < 0.0001 | < 0.0001 | < 0.0001 | < 0.0001 |
| Emotion Control | < 0.0001 | < 0.0001 | < 0.0001 | < 0.0001 | < 0.0001 |
| TD > ASD | GU Site | KK Site | Combined Sample | Split Half 1 | Split Half 2 |
| FIQ | - | < 0.0001 | 0.02 | - | 0.005 |
| **VIQ** | - | 0.001 | 0.031 | - | 0.006 |
| **PIQ** | - | 0.011 | 0.034 | - | 0.036 |

**Table 4.** Correlations between the three subscales of behavioral regulation, age, sex, motion, IQ and SRS. Results are given for the combined sample and as r-values of bivariate correlations. Motion (in mm) refers to the absolute maximum displacement at any timepoint in the resting-state fMRI scan prior to motion mitigation and denoising procedures. IQ=Intelligence Quotient; FIQ=full scale IQ; VIQ=verbal IQ; PIQ=performance IQ; SRS=Social Responsiveness Scale (total score). *denotes significance at p<0.05 uncorrected; ** denotes p<0.00055 (p<0.05 Bonferroni corrected for 90 comparisons).

|  | Age | Sex | Motion | FIQ | VIQ | PIQ | SRS | Inhibition | Shifting | Emotion Control |
| --- | --- | --- | --- | --- | --- | --- | --- | --- | --- | --- |
| Age |  | -0.19* | -0.17* | 0.05 | 0.04 | 0.03 | 0.03 | 0.11 | 0.05 | 0.01 |
| Sex | -0.19* |  | -0.07 | 0.02 | -0.05 | -0.05 | -0.07 | -0.01 | -0.09 | -0.05 |
| Motion | -0.17* | -0.07 |  | -0.18* | -0.10 | -0.17* | 0.17* | 0.14* | 0.10 | 0.11 |
| FIQ | 0.05 | 0.02 | -0.18* |  | 0.80** | 0.77** | -0.23** | -0.21** | -0.20* | -0.07 |
| VIQ | 0.04 | -0.05 | -0.10 | 0.80* |  | 0.44** | -0.18* | -0.13* | -0.11 | -0.01 |
| PIQ | 0.03 | -0.05 | -0.17* | 0.77** | 0.44** |  | -0.18* | -0.14* | -0.16* | -0.05 |
| SRS | 0.03 | -0.07 | 0.17* | -0.23** | -0.18* | -0.18* |  | 0.77** | 0.83** | 0.69** |
| Inhibition | 0.11 | -0.01 | 0.14* | -0.21** | -0.13* | -0.14* | 0.77** |  | 0.70** | 0.61** |
| Shifting | 0.05 | -0.09 | 0.10 | -0.20* | -0.11 | -0.16* | 0.83** | 0.70** |  | 0.79** |
| Emotion Control | 0.01 | -0.05 | 0.11 | -0.07 | -0.01 | -0.05 | 0.69** | 0.61** | 0.79** |  |

### 3.2 FC Models of Behavioral Regulation

Significant models were built using negative edges for inhibition (r_s_=.23, p=0.0001) and shifting (r_s_=.19, p=0.001), using leave-one-out prediction (N-1 at every iteration). Positive edge models were not significant for these measures and neither positive nor negative edge models were significant for emotion control (r_s_<.05, p>0.4). The FC model of inhibition was significant by permutation testing (r_s_=.23, p=0.037), while the FC model of shifting fell just shy of significance (r_s_=.19, p=0.067). As seen in Figure 1, inhibition revolved around edges in the somato-motor, visual and cerebellar networks and was more posterior, while shifting appeared more focused on edges around the frontoparietal and dorsal attention networks and was more anterior. Both inhibition and shifting included a number of edges connecting with DMN regions as well as the temporal lobe. Note that in the leftmost panels higher rank refers to a lower score, i.e. lower symptoms. In negative edge models, lower FC associates with higher ranked scores.

**Figure 1.**
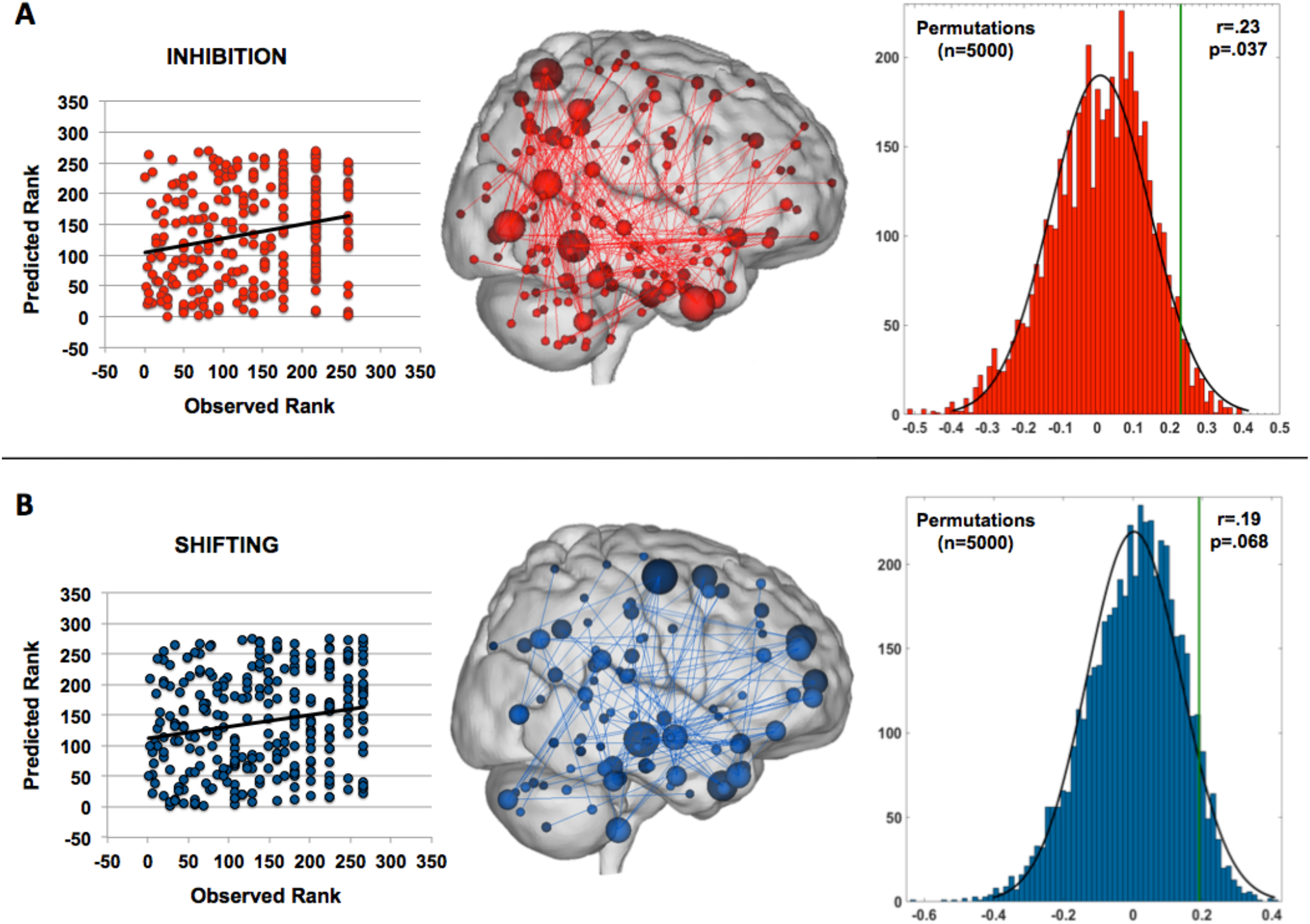
CPM models for inhibition (panel A) and shifting (panel B). Models are evaluated using a leave-one-out approach, with a different participant left out in each iteration. The predictive potential is assessed by comparison of the predicted and actual score ranks (left column; inhibition: r=.23; shifting: r=.19) using Spearman’s rank correlation, and statistical significance for the correlation between predicted and observed values is assessed using permutation testing (right column; inhibition: p=0.037; shifting: p=0.068). The inhibition model revolved around edges in the somato-motor, visual and cerebellar networks (upper middle column) and was more posterior/inferior, while shifting appeared more focused on edges around the frontoparietal and dorsal attention networks (lower middle column) and was more anterior. Both inhibition and shifting included a number of edges in the default mode network (DMN) and the temporal lobe. The size of the nodes reflects the number of connections the node has to other nodes, with larger nodes being more connected than smaller nodes.

### 3.3 Cross-Validation: Split Halves Prediction

Significant models were built for inhibition and shifting behavioural regulation measures using the negative edges in a leave-one-out prediction (N-1 at every iteration) in both split half 1 (inhibition: r_s_ =.26, p=0.002 and shifting: r_s_ =.32, p=0.0001) and split half 2 (inhibition: r_s_=.17, p=0.049 and shifting: r_s_=.34, p=0.00005). The models built in split half 1 further significantly predicted scores in the unseen second half (inhibition: r_s_=.39, p<0.000001 and shifting: r_s_=.19, p=0.03), and the models built in split half 2 significantly predicted scores in the unseen first half (inhibition (r_s_=.48, p<0.000001 and shifting (r_s_=.19, p=0.02).

### 3.4 Cross-Validation: From Site 1 to Site 2 Prediction

Models could not be built for shifting or inhibition behavioural regulation measures using the negative edges in a leave-one-out prediction (N-1 at every iteration) in the GU site (inhibition: r_s_=.15, p=0.17 and shifting: r_s_=.09, p=0.4) or in the KK site (inhibition (r_s_=.02, p=0.8) and shifting (r_s_=.03, p=0.71). Therefore, no cross-prediction from GU to KK and vice versa was attempted.

### 3.5 Model Specificity to Behavioral Regulation

In the combined model, Spearman rank correlations between observed and predicted score ranks for TD children (n=191) and children with ASD (n=85) separately were insignificant for the smaller ASD group in both inhibition (r_s_=.14, p=0.22) and shifting (r_s_=.10, p=0.36), but near significant in the TD group in inhibition (r_s_=.14, p=0.053) and significant in shifting (r_s_=.22, p=0.002) (Figure 2). Spearman correlations between predicted shifting or inhibition and motion or IQ were insignificant (r_s_’s<.10). Both predicted inhibition (r_s_=.16, p=0.01) and shifting (r_s_=.23, p<0.001) scores associated significantly with total SRS scores; this association fell just shy of significance in a partial correlation when controlling for diagnosis (inhibition and SRS r_s_=.11, p=0.069; shifting and SRS r_s_=.12, p<0.051). In addition, predicted shifting scores significantly associated with observed emotional control scores (r_s_=.14, p=0.018), but not inhibition (r_s_=.06, p=0.33). Similarly, predicted inhibition scores weakly associated with observed shifting scores (r_s_=.12, p=0.04), but not emotional control scores (r_s_=.1, p=0.09). Neither shifting nor inhibition significantly associated with motion, neither before cleaning (r_s_<0.05) nor after (r_s_<.11). Note all associations reported in this section are uncorrected for the number of comparisons performed.

**Figure 2.**
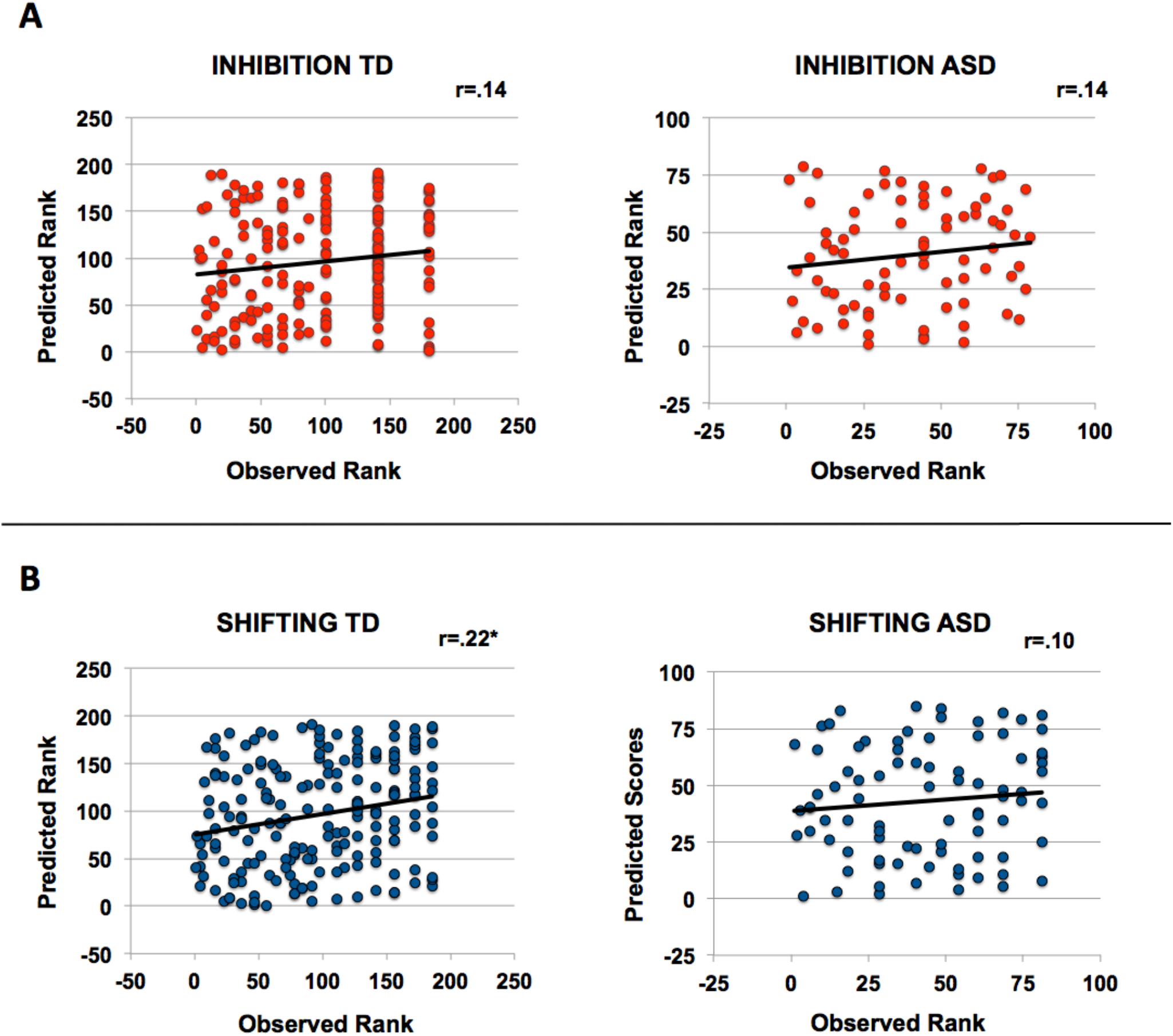
Spearman rank-correlations between observed and predicted score ranks of inhibition (panel A) and shifting (panel B) for TD children and children with ASD, separately to evaluate whether our models capture behavioral regulation dimensionally or are driven by the categorical difference in scores due to ASD diagnoses. Results were insignificant for the smaller ASD group in both inhibition and shifting, but near significant in the TD group in inhibition and significant in shifting.

## 4 DISCUSSION

Children with ASD are known to struggle with behavioral self-regulation, which associates with greater daily-life challenges and an increased risk for psychiatric comorbidities [11, 13, 67, 68]. In a fully cross-validated, data-driven analysis in a large sample of children with and without ASD that was compiled across two data collection sites, we identified distributed patterns of FC whose strength predicted individual differences in two behavioral regulation subdomains. These whole-brain network models predicted novel individuals’ inhibition and shifting, but not emotional control scores from resting-state FC data both in a leave-one-out, as well as in a split halves cross-validation, providing evidence that meaningful correlates of behavioral regulation in intrinsic brain patterns exist. Indicating the limitations of this approach, whole-brain network models could, however, not be built within the smaller and less balanced samples collected at the respective sites. We further found that although models captured within-group variation, models built on inhibition and shifting also predicted ASD symptoms more generally, and this relationship persisted at a trend-level after taking diagnosis into account. Overall our results, showing that complex brain network models predict different measures of behavioral regulation across a sample of children with and without ASD, demonstrate that whole-brain FC data can serve as a holistic neural index of inhibition and shifting.

Our findings present a substantial addition to our knowledge on the neural expressions of inhibition and shifting across the spectrum of children with and without ASD. The majority of neuroimaging research on inhibition and shifting has been done in neurotypical adults, and little is known about neural alterations underlying shifting and inhibition for children with ASD. Encouragingly, our findings are broadly in line with existing literature. Both alterations in inhibition and shifting have previously been associated with changes in the DMN - which has hubs in anterior cingulate and ventromedial PFC - in a number of disorders. For instance, activity in regions of the DMN has been shown to be altered in relation to inhibition in ADHD (in a stop signal paradigm [69]), OCD (during a reward paradigm [70]), and major depression (in a cognitive control paradigm [71]). Similarly, the DMN has been shown to be altered in relation to shifting in Schizophrenia (during reinforcement learning [72]), OCD [73] and ADHD [74]. In ASD, alterations in FC involving DMN regions have been linked to inhibition [19] and SRS scores [75], which correlate with both inhibition and shifting in our sample.

The divergent neural underpinnings of inhibition and shifting shed further light into children’s functional brain mechanisms. The cerebellum is heavily implicated in ASD [76–78] and our group has previously observed that cerebellar FC related to ADHD hyperactivity scores in young children aged 4-7 years old [79], which provide an index of inhibition [80, 81]. Likewise, somatosensory sensitivities in ASD – like repetitive motor or tactile behaviors etc. – are known to correlate with neural alterations in somato-motor areas [82], and the ability to inhibit such behaviors is linked to inhibition [2]. Individuals with ASD also manifest anomalies in their visual selection and have greater difficulty than neurotypical populations when ignoring specific visual inputs, which relate to alterations in the visual stream [83]. During set-shifting tasks, increased activation in parietal lobes has been reported in individuals with ASD [1], and our group has previously observed that FC in the dorsal attention network related to ASD attention switching scores in young children [79].

Our work further evidences two major challenges that remain towards achieving one of the primary goals of human neuroimaging - to identify generalizable neuromarkers of clinical utility. First, whole-brain network models could not be built successfully within the smaller and less balanced samples collected at the respective sites. It is a known problem in ASD research that there are site effects that appear to influence generalizability, meaning that it may be unlikely to build generalizable predictive markers using data from a single site. Second, models built on inhibition and shifting scores also predicted ASD symptoms more generally. While this is to be expected given the known association between ASD core social symptoms and behavioral regulation, it is worthwhile to note that since these relationships persisted at trend-level when diagnosis was controlled for, they are unlikely to be entirely driven by diagnostic category. In addition, relationships between observed and predicted shifting scores remained significant when assessed only within the TD population, again indicating that the predictions have some specificity to the constructs under investigation. Nonetheless, the correlation between social and behavioral regulation traits makes it more difficult to tease apart the unique and shared features between these constructs.

Individual differences in relation to behavioral regulation have been repeatedly found to be associated with individual features in FC [33, 39, 84]. Taking individual differences into account can help expose the underlying neural substrates of complex cognitive skills, emotions, social competencies and more, and has proven useful in the investigation of both neurotypical [38, 41, 85, 86] and clinical populations [87–89]. It has been argued that clinical cut-offs for diagnosis may be arbitrary [45, 87], as traits and abilities associated with neurodevelopmental disorders such as ASD also exist in the neurotypical population, and fall onto a spectrum or continuum [87, 90]. The use of dimensional approaches also allows for more statistical power in studies on neurodevelopmental disorders, which are chronically underpowered due to small sample sizes and challenged by heterogeneity in the populations studied [45, 91–93]. Current research suggests that neural patterns associated with abilities that are affected by neurodevelopmental disorders are both categorical (i.e. unique to a diagnosis) as well as dimensional (i.e. also present in TD populations) [94, 95]. Our findings point to distinct neural mechanisms in the brain subserving the different subtypes of behavioral regulation, which may aid in informing us about options for targeted interventions. They thus highlight possibilities for gleaning insight into how the brain’s functional organization may be associated with cognitive and behavioral issues in children and may serve as a basis for future studies investigating behavioral regulation in neurodevelopmental and other disorders.

The current study has several distinct strengths, which include the use of three measures of behavioral regulation in two relatively large, “matched” groups of children with and without ASD and novel preprocessing techniques. Both samples were acquired in close spatial and temporal proximity, that is, around the same time and around the same geographic location (Washington, D.C. and Baltimore). The assessment of behavioral regulation is well validated [16] and although parent- and self-reports are subjective, they capture a measure of behavior integrated over a longer time frame than can be observed in a laboratory visit and have better test-retest reliability [96]. At both sites, children were introduced to an MRI simulator first and given the opportunity to familiarize themselves with the experience of undergoing an MRI scan with their eyes open. The denoising methods employed allow for the retention of the remaining ‘true’ neural signal within an affected volume and are in accordance with the latest best practices for reducing motion artifacts [59], which is particularly relevant for a neurodevelopmental study [57, 97, 98].

The study also has several weaknesses, including differences within and between the GU and KK sites. These are differences in (i) fMRI data acquisition procedures, (ii) in-/exclusion criteria, (iii) numbers of participants, (iv) TD vs. ASD ratios, (v) sex ratios, (vi) IQs, (vii) BRIEF scores, (viii) comorbidities, and (ix) medications. All of these serve as a reminder of just how divergent ASD populations are, how difficult it is to assess whether a sample reflects the “true patient population” and to what extent research findings are truly generalizable. Considering that these two sites contain a notable number of participants also highlights how common datasets with unbalanced groups are – a known problem when performing cross-validation and assessing prediction performance [65]. Another weakness potentially rests in the removal of motion-associated edges where motion correlates with the behavior that is investigated. While in the combined sample, motion did not significantly associate with the behavioral regulation measures after multiple comparison correction, there remains a possibility that edges that would have associated to, and strengthened the behavioral regulation models, were removed. The same holds true for site-specific associations. Finally, emotional control scores were not significantly predicted for novel individuals. It is perhaps unsurprising that emotional control results did not follow the same pattern as the other measures of behavioral regulation, as evidence suggests that definitions of emotional control abilities are broad, difficult to dissociate from emotion generation processes, and fairly hard to capture reliably [25, 99–101]. Another possibility is that the neural mechanisms of emotional control are simply not reflected in whole-brain functional connectivity patterns consistently across individuals with ASD.

It should be noted that like most methodological approaches, CPM has both advantages and challenges. One major advantage is that because CPM models are defined and validated with independent data, they promise to improve our ability to uncover generalizable brain-behavior associations [65, 102]. A major challenge is that predictive models based on FC will only ever account for a fraction of the variance, because they are limited by how much information the signal can capture as well as the chosen phenotypic measure. They are also bound by linearity assumptions: Linear models built across TD children and children with ASD can capture dimensional associations, but may miss categorical differences, which are distinct from dimensional associations (please see [94, 95] for a discussion on this). Finally, one may consider the current CPM framework as perhaps a bit simplistic in that it yields only one summary value for ‘positive’ and ‘negative’ networks, cannot capture flexible brain network dynamics, and has no ‘blueprint’ for how to tie together predictions on a multitude of behavioral measures. With this being said, as Scheinost and colleagues [65] have so eloquently put it: *“When studying brain-behavior associations, one must keep in mind how extraordinary it is that neuroimaging data can be distilled to approximate phenotypic measures that reflect a simplification of multiple complex features. Thus, even modest results are reasonable and remarkable”*.

Future studies could benefit from using data that was obtained while participants perform a task or are under naturalistic viewing conditions such as movie watching. Data obtained while participants perform a task, which adequately captures differences in abilities or skills, has been shown to associate with differences in FC and to lead to better predictive models [62, 103]. Showing videos increases young children’s ability to stay still during a scan [104], and may be especially useful for studies in children with neurodevelopmental disorders, many of whom evidence attention difficulties in addition to challenges staying still for MRI acquisitions [63, 89, 105–109]. It was further recently shown that individual differences in FC are enhanced during passive viewing, thus facilitating their detection not only through reduced motion but also through the synchronization of hemodynamic fluctuations in large areas of the cortex across participants itself [110]. Another improvement could be yielded through the implementation of longer scan imaging times to strengthen the reliability of FC estimates, allow for some data loss in wiggly children and the use of other existing motion mitigation techniques (see e.g. [79, 111, 112]. Finally, harmonized in- and exclusion criteria and scanner and experimental protocols could also aid in providing more comparable FC estimates [47, 113].

The characterization of differential presentation of ASD core symptoms and ASD associated symptoms along the spectrum has been of immense interest to researchers, as heterogeneity is a defining characteristic of ASD that makes it difficult to assess and treat within traditional diagnostic frameworks. In this study, we utilized a data-driven approach to develop objective quantitative FC models that elucidate and predict performance in behavioral regulation subdomains in ASD. We observed both commonalities and differences in the functional organization of inhibition and shifting across TD children and children with ASD, with inhibition relying on more posterior and shifting relying on more anterior brain networks. We also demonstrate the generalizability and trans-diagnostic utility of this approach, as well as its clear limits to date. Given the numerous cognitive and behavioral issues that can be quantified dimensionally in neurodevelopmental disorders - and literally any disorder or disease - further refinement of whole-brain neuromarker techniques will prove vital to neuroimaging’s role in individualized medicine.

## DECLARATIONS

### Abbreviations

ABIDE-II: Autism Brain Imaging Data Exchange II
ASD: Autism Spectrum Disorder
BRIEF: Behavior Rating Inventory of Executive Function
CPM: Connectome Predictive Modelling
DMN: Default Mode Network
FIQ: Full-scale IQ
FC: Functional Correlation or Functional Connectivity
GU: Georgetown University
KK: Kennedy Krieger Institute
PIQ: Performance IQ
SRS: Social Responsiveness Scale
TD: Typically Developing
VIQ: Verbal IQ
WISC-IV: Wechsler Intelligence Scale for Children - Fourth Edition
WISC-V: Wechsler Intelligence Scale for Children - Fifth Edition
WASI: Wechsler Abbreviated Scale of Intelligence

## Acknowledgements

**Acknowledgements**

## Availability of Data and Materials

The datasets analyzed in the current study are publicly available under fcon_1000.projects.nitrc.org/indi/abide/abide_II.html. CPM scripts are publicly available at www.nitrc.org/projects/bioimagesuite.

## Authors’ Contributions

CR and SB designed the study and analyzed and interpreted the data. SK made substantial contributions to the analysis through adaptation of scripts and computational testing. CR wrote the manuscript with input from SB. All authors read and approved the final manuscript.

